# DNA methylation of transposons pattern aging differences across a diverse cohort of dogs from the Dog Aging Project

**DOI:** 10.1101/2024.10.08.617286

**Authors:** Blaise L. Mariner, Brianah M. McCoy, Ashlee Greenier, Layla Brassington, Elizabeth Slikas, Christine Adjangba, Abbey Marye, Benjamin R. Harrison, Tal Bamberger, Yadid Algavi, Efrat Muller, Adam Harris, Emily Rout, The Dog Aging Project Consortium, Anne Avery, Elhanan Borenstein, Daniel Promislow, Noah Snyder-Mackler

**Author notes:** equal contribution (Blaise L. Mariner, Brianah M. McCoy). **Statements and declarations None**. No competing interests.

## Abstract

Within a species, larger individuals often have shorter lives and higher rates of age-related disease. Despite this well-known link, we still know little about underlying age-related epigenetic differences, which could help us better understand inter-individual variation in aging and the etiology, onset, and progression of age-associated disease. Dogs exhibit this negative correlation between size, health, and longevity and thus represent an excellent system in which to test the underlying mechanisms. Here, we quantified genome-wide DNA methylation in a cohort of 864 dogs in the Dog Aging Project. Age strongly patterned the dog epigenome, with the majority (66% of age-associated loci) of regions associating age-related loss of methylation. These age effects were non-randomly distributed in the genome and differed depending on genomic context. We found the LINE1 (long interspersed elements) class of TEs (transposable elements) were the most frequently hypomethylated with age (FDR < 0.05, 40% of all LINE1 regions). This LINE1 pattern differed in magnitude across breeds of different sizes– the largest dogs lost 0.26% more LINE1 methylation per year than the smallest dogs. This suggests that epigenetic regulation of TEs, particularly LINE1s, may contribute to accelerated age and disease phenotypes within a species. Since our study focused on the methylome of immune cells, we looked at LINE1 methylation changes in golden retrievers, a breed highly susceptible to hematopoietic cancers, and found they have accelerated age-related LINE1 hypomethylation compared to other breeds. We also found many of the LINE1s hypomethylated with age are located on the X chromosome and are, when considering X chromosome inactivation, counter-intuitively more methylated in males. These results have revealed the demethylation of LINE1 transposons as a potential driver of intra-species, demographic-dependent aging variation.

## Introduction

Life expectancy is generally inversely correlated with body size within mammalian species [1,2], while females typically live longer than males [3]. Despite the various mechanisms influencing these (and other [4,5]) demographic differences in lifespan, we still know little about how epigenetic differences between individual characteristics may also contribute to the variation in lifespan, health, and disease within a mammalian species. Companion dogs (*Canis familiaris*) are a powerful translational model with immediate relevance to human health [6], exemplified by their great genetic and phenotypic diversity, dynamic and variable environments, diverse diets, well-established healthcare system, and wide geographical distribution— many of which are characteristics shared with humans [7–10].

<u>T</u>ransposable <u>e</u>lements (TEs) are selfish genetic elements that propagate by transposing (or “jumping”) around the genome [11–13]. TE transposition can cause genomic instability and contribute to the onset of cancers and other age-related diseases [14–23]. TE activity is generally suppressed by epigenetic modifiers such as <u>DNA</u> <u>m</u>ethylation (DNAm), a mechanism organisms generally use to control cell fate and function [24,25]. Changes in DNAm patterns close to TEs can have wide-ranging consequences [26–31], but largely, the loss of methylation (hypomethylation) at TEs specifically is a known consequence of aging [25,32–34]. Interventions that suppress TE transposition can extend the lifespan in *Drosophila* and *Caenorhabditis elegans*, suggesting that these genetic parasites play a role in regulating aging [35–38]. In mice, dysregulation of TEs can lead to accelerated aging and is partially rescued with drugs that can repress TE transposition [39–41]. Research aiming to mitigate TE-related genomic damage has wide-ranging (e.g. oncogenic) practical applications [11–13,42,43]. Individual characteristics (e.g. size, sex) of dogs may influence TE regulation, which may help explain the variation in aging and disease onset between individuals of the same species.

Given that dogs exhibit notable variation in aging rates, with body size being a well-known factor influencing this variation [2,44,45], the regulation of TEs may offer additional insights into aging and disease onset within a species [46]. DNA methylation (DNAm) at TEs, known to suppress TE activity, could serve as a useful proxy for investigating how TE activity correlates with individual characteristics such as size and sex. By examining whether DNAm patterns near TEs differ according to these characteristics, we can begin to explore whether variation in TE regulation might help explain differences in the rates of aging and disease onset among dogs [32].

Here, we generated genome-wide DNAm data from immune cells of 864 dogs as part of the Dog Aging Project (DAP) to characterize age-related 1) global DNAm changes, 2) TE-specific changes, and 3) demographic drivers of these differences in 1 and 2. Based on the existing literature [32,43,47,48], we hypothesized that the methylation of TE regions would decrease with age in dogs. We found that not only are TEs hypomethylated with age, but larger dog breeds have more accelerated loss of TE methylation. We also found that TEs hypomethylated with age are overwhelmingly located on the X chromosome and are, on average at all ages, more methylated in males– a surprising result counter-intuitive to X chromosome inactivation.

## Results

### The aging methylome of canine immune cells

We measured DNA methylation of CpG sites in peripheral blood mononuclear cells (PBMCs) collected from 864 dogs (420 female, 444 male) of diverse breed sizes (195 Small, 183 Medium, 199 Standard, 215 Large, 72 Giant) using reduced-representation bisulfite sequencing (RRBS) (**Figure 1, Supplemental Figure 1A & 1B**, see methods). We quantified DNAm for 3,063,134 well-covered CpG sites across the dog genome, which we combined into 47,393 CpG regions to reduce the number of hypothesis tests and the amount of missing data due to stochastic variation in sequencing coverage (see methods and [49,50]). We then tested if age or breed size predicted CpG region DNAm in a binomial mixed-effects model controlling for other covariates and genetic relatedness [51].

**Figure 1:**
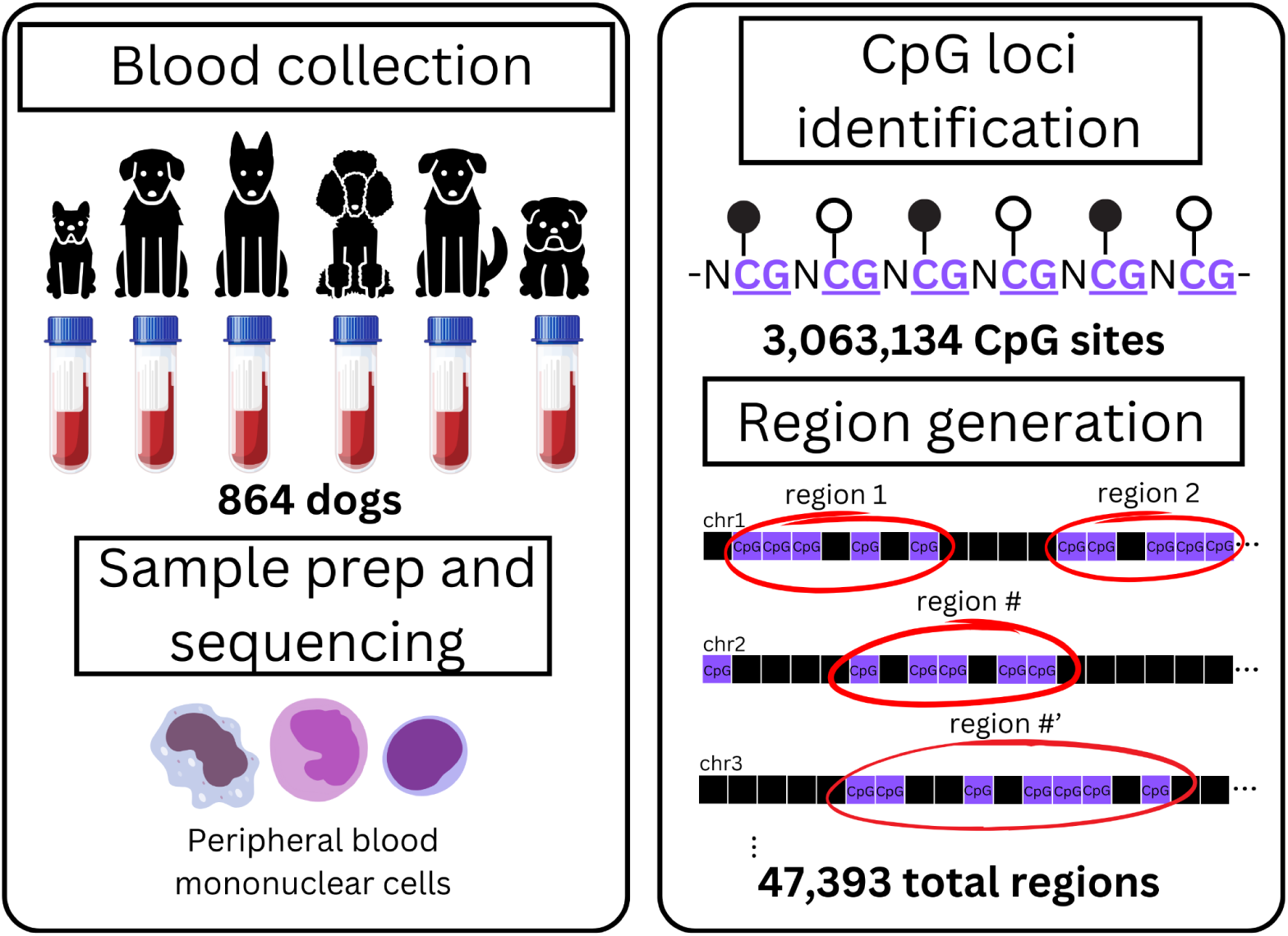
Sample workflow and region generation. Veterinarians collected blood samples from Dog Aging Project (DAP) dogs. Researchers then isolated and prepped peripheral blood mononuclear cells (PBMCs) for reduced-representation bisulfite sequencing (RRBS) to detect 5’-cytosine-phosphate-guanine-3’ (CpG) methylation. CpG loci were grouped into genomic regions and annotated using their genomic coordinates (methods).

Age strongly patterned DNAm across the genome: 42% of the tested regions were significantly associated with age (FDR<0.05). The majority (66%) of these regions were hypomethylated in older dogs compared to younger dogs (**Figure 2A**), consistent with the known age-associated global loss of methylation observed in many other species [52–54].

**Figure 2:**
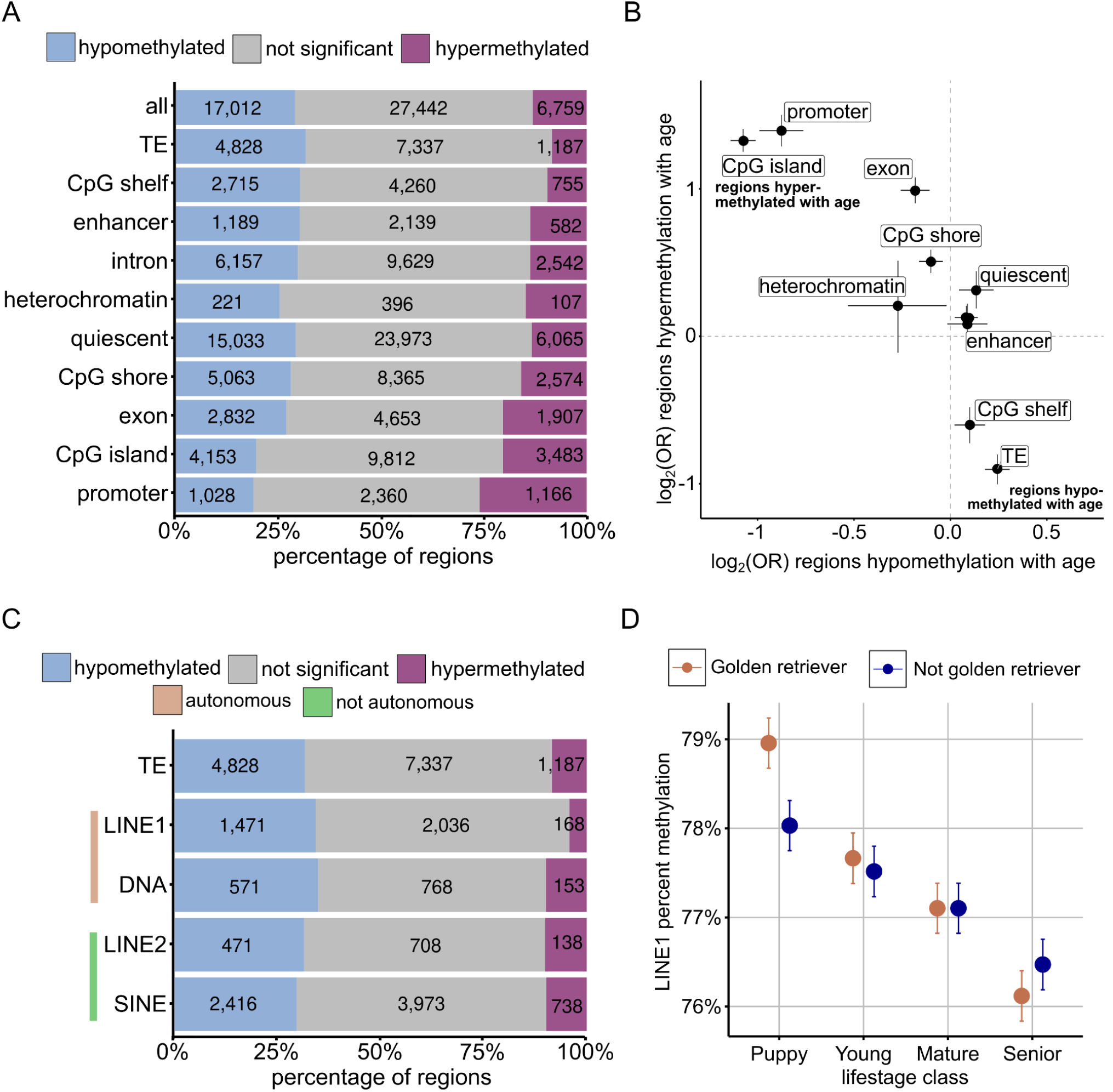
Age differentially affects DNA methylation across the genome. A) Percent of regions in each annotation category hypomethylated with age (blue; β_age_ < 0 & FDR < 0.05), hypermethylated with age (purple; β_age_ > 0 & FDR < 0.05), or not associated with age (grey). B) Enrichment (log_2_OR) of regions hypomethylated (x-axis; done with regions β_age_ < 0 & FDR < 0.05) and hypermethylated (y-axis; done with regions β_age_ > 0 & FDR < 0.05) with age. Error bars: 95% confidence intervals. C) Percent of regions in each TE class hypomethylated with age (blue; β_age_ < 0 & FDR < 0.05), hypermethylated with age (purple; β_age_ > 0 & FDR < 0.05), or not associated with age (grey). D) Mean percent methylation of LINE1s by life stage classification of golden retrievers (brown) and non-golden retrievers (blue).

Age-associated CpG regions were non-randomly distributed in the genome–likely reflecting context-dependent effects of age on DNAm. Specifically, compared to all age-associated regions, promoters were twice as likely to be hypermethylated with age (1,115 regions; 26% of all promoter regions; log_2_(OR) = 1.39, p_adj_ < 2e-16) (**Figure 2B**). By the same measure, we also found that exons (log_2_(OR) = 0.99, p_adj_ < 2e-16), CpG islands (which are CpG-rich regions in the genome) (log_2_(OR) = 1.32, p_adj_ < 2e-16), and CpG shores, which flank CpG islands, (log_2_(OR) = 0.51, p_adj_ < 2e-16) were more likely to be hypermethylated with age compared to the background expectation across all regions. Conversely, we found TEs and CpG shelves, which flank CpG shores, were significantly under-represented in the regions hypermethylated with age (TEs: log_2_(OR) = −0.90, p_adj_ < 2e-16; CpG shelves: log_2_(OR) = −0.60, p_adj_ < 2e-16) and over-represented in the regions hypomethylated with age (TEs: log_2_(OR) = 0.24, p_adj_ < 2e-16; CpG shelves: log_2_(OR) = 0.10, p_adj_ = 0.02). This age-associated loss of methylation in TEs and CpG shelves may reflect these regions’ increased activity–e.g., TE transposition and/or distal gene regulatory (e.g. enhancer) activity.

### Functional enrichment of genes with age-associated promoter DNAm

Genes with promoters hypermethylated with age were enriched for bone marrow and blood cell markers (**Supplemental Figure 3A**), likely reflecting the changing dynamics of these immune cells’ gene regulation with age. Additionally, these genes were also more likely to be targets of PRC2 (<u>p</u>olycomb <u>r</u>epressive <u>c</u>omplex <u>2</u>) and H3K27me3–both of which have been previously linked to age-associated DNAm patterns (**Supplemental Figure 3B**) [55,56]. Both PRC2 and H3K27me3 repress TE activity, and thus, age-related hypermethylation of their binding motifs may reflect a growing inability to repress TEs with age [55,57,58]. We also found that genes differentially methylated with age were associated with developmental pathways and cancer signatures. This cancer-related signature is concentrated in promoters with age-associated hypermethylation, which were enriched for genes that are up-regulated in response to tretinoin treatment (a drug used to treat leukemia [59]) and genes implicated in lung and liver cancers [60,61].

Using motif enrichment analysis, we further tested the sequences of the 2kb promoter regions to uncover transcription factor binding motifs differentially methylated with age. We found that promoters hypermethylated with age were enriched for numerous motifs, including enrichment of transcription factors involved in embryo development, immune response, inflammation, and stress response signaling (**Supplemental Figures 3C-3E**). For example, the <u>a</u>ctivating transcription factor <u>4</u> (ATF4) binding motif (5’-GTGACGT[AC][AG]-3’) is one of the stress response motifs hypermethylated with age [62]. ATF4 is known to be up-regulated in many types of long-lived mice, and ATF4’s orthologs are necessary for lifespan extension in several different aging model organisms [63–67]. Hypermethylation of transcription factor motifs would likely impact transcription factor binding and downstream activity influencing gene expression.

### Autonomous TEs become less methylated with age

TEs, which have been implicated in aging across many species, had the strongest signal of age-associated hypomethylation–likely reflecting that they may become more active with age. However, “transposable elements” is an umbrella term that is comprised of various types, or “classes”, that differentially impact the host. Briefly, TEs have different mechanisms for transposition–some require RNA intermediaries (e.g., LINEs and SINEs), and others solely DNA (e.g., DNA transposons [68]). TEs can also be broadly categorized into “autonomous” and “non-autonomous” classes, where autonomous TEs can transpose without any help from the host. In contrast, non-autonomous host sequences or machinery can mobilize [69]. The autonomous class consists of (1) DNA transposons [68] and (2) LINE1 (long <u>in</u>terspersed <u>e</u>lements) that carry their own reverse-transcription machinery. LINE2s and SINEs (<u>s</u>hort <u>in</u>terspersed <u>e</u>lements), however, are non-autonomous and require other elements’ reverse-transcription machinery to transpose. We thus carried out a more nuance analysis to identify which classes may be driving the observed age-associated patterns in dogs.

Autonomous TEs were the most frequently hypomethylated with age, with significant age-associated loss of methylation in 34% of all LINE1s (n = 3,305) and 34% of all DNA transposons (n = 1,388). In contrast, 31% of LINE2s (n=1,209) and 30% of the SINEs (n=6,555) were hypomethylated with age (**Figure 2C**). Of all the TE classes, LINE1s showed the lowest amount of hypermethylation with age: only 4% of LINE1s were significantly hypermethylated with age compared to ∼10% in all other TE classes (**Figure 2C**). On average, all TE classes became less methylated with age, but this decline was the steepest in LINE1s (β_LINE1_ = −0.13%; β_DNATransposon_ = −0.12e-3%; β_LINE2_ = −0.10%; β_SINE_ = −0.089%). When we compared this decline in TEs relative to the whole genome, we similarly found the fastest decline in LINE1s (and DNA transposons) methylation relative to the entire genome (β_LINE1_ = −5.57e-2%, p = 5.7e-9; β_DNATransposon_ = −5.4e-2%, p < 2.2e-16; β_LINE2_ = −3.8e-2%, p < 2.2e-16; β_SINE_ = −2.6e-2%, p < 5.6e-15; similar to [32]). This suggests that age-associated loss of methylation in LINE1s, and to a lesser extent DNA transposons, outpaces other TE classes’ age-associated methylation loss.

Given the strong age-associated hypomethylation in LINE1s and the link between LINE1 demethylation and cancer onset [70], it is possible that variation in this age-associated loss of methylation at LINE1s might reflect variation in cancer incidence across breeds. To test this thoroughly, however, requires precise pairing of breed-specific cancer prevalence with breed assignments of our study animals–data that are outside of the scope of this manuscript. Nevertheless, we attempted a preliminary test of this link between LINE1 demethylation and cancer risk in golden retrievers, the most common breed (n=44) in our study, which also have an extremely high risk of hematopoietic cancer ([31,71–74] note that our data were generated in immune cells). Interestingly, we found that golden retrievers have faster age-related methylation loss of LINE1 elements than non-golden retrievers (β_age:breedGR_ = 6.67e-2%, p = 0.585; **Figure 2D**). This result suggests that increased TE activity in older golden retrievers may be mechanistically linked to their increased risk of hematopoietic cancers, a disease state whose cause has remained elusive to veterinarians [75].

### X-chromosome is enriched for age-associated LINE1s

LINE1 activity and impact varies across chromosomes, and it is well-established that some chromosomes, particularly the X chromosome, are particularly enriched for LINE1s [69,76,77]. Thus, we next sought to characterize the chromosomal distribution of age-associated LINE1s, which, if biased, might provide mechanistic insight into sex differences in aging and mortality. Of the 1,130 LINE1s hypomethylated with age (FDR < 0.05), 245 of them are located on the X chromosome (log_2_(OR) = 3.1, p_adj_ < 2e-16) (**Figure 3A**). We hypothesized that these X-linked LINE1s would be more methylated in females than males–in large part due to X chromosome inactivation in females where one copy of the X chromosome is almost completely hypermethylated. To test this, we first compared mean percent methylation between males and females of all age-associated X-linked non-LINE1 regions. As we would expect, DNAm of these regions was significantly higher in females compared to males (β_Male_ = −3.5% ; p<2e-16). However, this pattern was reversed when we compared DNAm only on X-linked LINE1s, which were more methylated in males (β_Male_= 0.4%, p = 3e-4; **Figure 3B**). This lower DNAm of X-linked LINE1s in females may correspond to increased LINE1 activity in females relative to males. This sex difference in X-linked LINE1 DNAm was ∼3x stronger when we limited our comparison to those LINE1 elements that were also significantly associated with age (β_Male_= 1.5%, p < 2e-16; **Figure 3C**). Together, these findings suggest that X-linked LINE1s are less methylated in females and thus cross a pathogenic threshold of activity earlier in females than in males.

**Figure 3:**
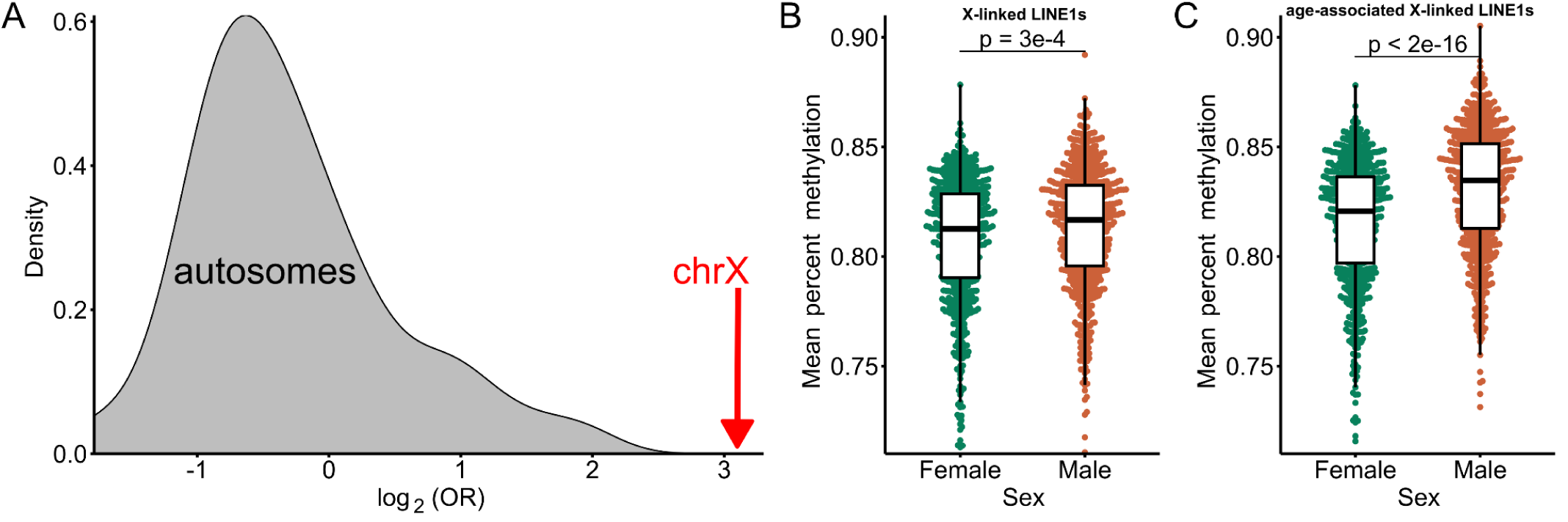
Chromosome locations of LINE1s hypomethylated with age. A) Density plot representing the origin of LINE1s hypomethylated with age by OR enrichment of all autosomes (grey) and the X chromosome (red). B) Mean percent methylation of all LINE1s on the X chromosome. C) Mean percent methylation of LINE1s on the X chromosome significantly associated with age (FDR < 0.05). P-values are from linear models: LINE1 percent methylation ∼ sex + age).

### Larger dogs show the strongest age-related DNAm changes

We next tested if dog size, which is negatively correlated with lifespan, predicted differences in the rate of DNAm changes with age. We used genetic predictions of adult dog height (**Supplemental Figure 1A & 1B**, [78]) to capture dog size rather than relying on breed averages or owner-reported weight. However, the results reported below were largely consistent when we used owner-reported and breed size estimates.

The effects of dog size on DNAm were significantly correlated with those of age on DNAm (Pearson’s r = 0.18, p < 2e-16), such that increasing size and age exerted similar overall effects on DNAm. After FDR correction, dog size was significantly associated with DNAm in 375 regions (FDR <0.05; note that this doubles to 819 at FDR < 0.1), which is only a fraction (1.5%) of the regions significantly associated with age. Nevertheless, these size-associated regions were enriched for age-associated regions, being twice as likely to also be age-associated than expected by chance (binomial test, p = 2.1e-7).

Given the significant overlap of age and size effects in LINE1 regions and the fact that LINE1s show the strongest age-associated loss of methylation, we next tested if larger dogs showed even faster rates of age-associated hypomethylation in these regions than other dogs. Indeed, age-associated LINE1 de-methylation was faster in larger dogs (β_Age:dog size_ = −0.37%, p = 0.013). This pattern was also evident when we categorized dogs into breed class sizes (**Figure 4**). On average, LINE1 elements become demethylated at a dramatically increasing rate from small dogs to giant dogs (β_Age:small dogs_ = +0.08% per year; β_Age:medium dogs_ = −0.08% per year; β_Age:standard dogs_ = −0.16% per year; β_Age:large dogs_ = −0.15% per year; β_Age:giant dogs_ = −0.24% per year). In short, giant dogs lose 0.26% more LINE1 methylation per year than the small dogs (β_Age:breedsizeGiant_ = −0.26%, p = 3.6e-3).

**Figure 4:**
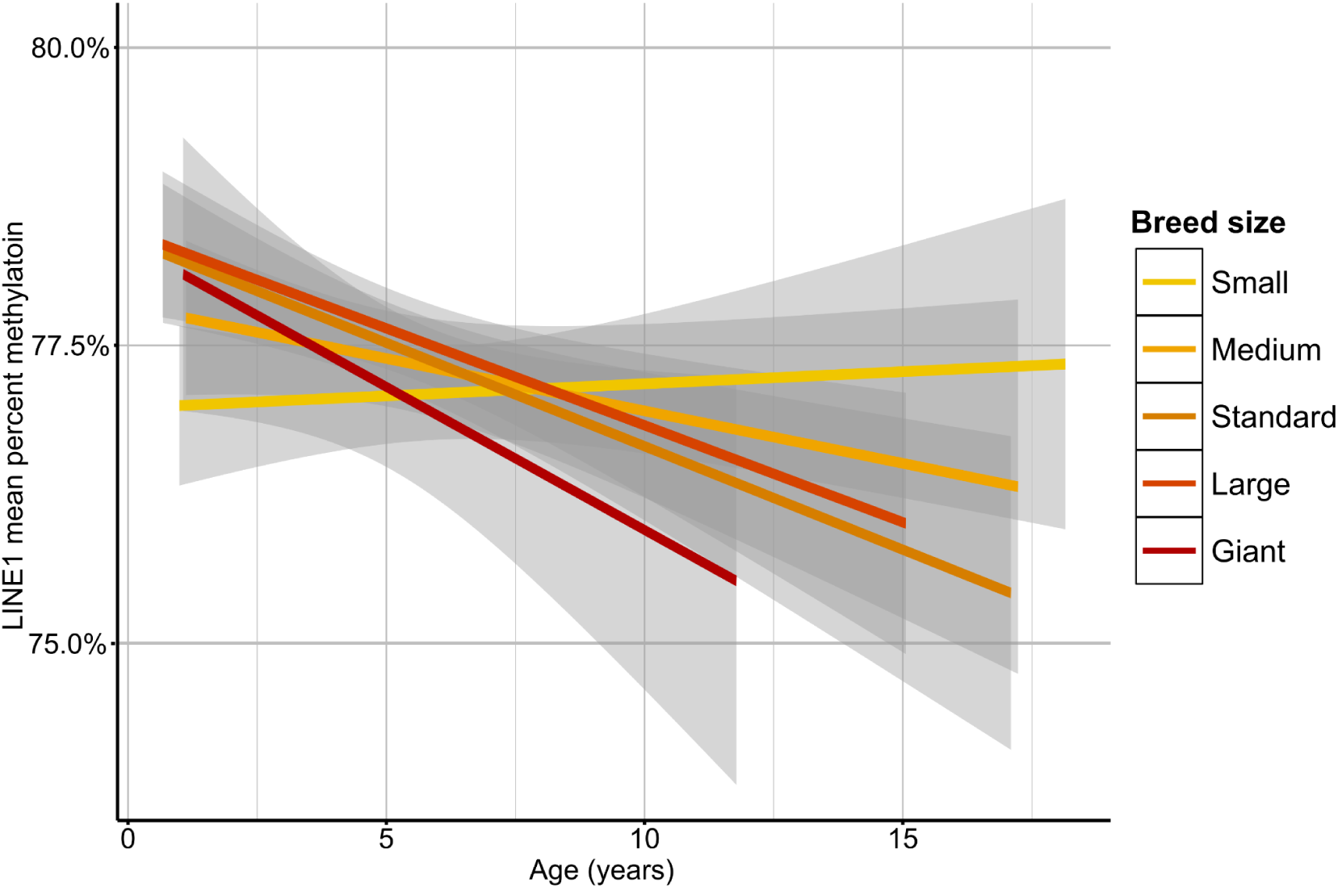
Breed size effects on age-related epigenetic acceleration. Mean percent methylation of all LINE1s by age across breed size classifications (linear model: LINE1 percent methylation modeled by Age:Breed size shown here for each breed size respectively). Shaded region represents the 95% confidence level interval for each nested effect respectively.

## Discussion

In this study we comprehensively examined the aging methylome of a large and diverse cohort of dogs from the DAP. We integrated the rich demographic data from the DAP with measures of DNAm across the dog genome to identify not only the normative molecular changes that occur with age in the dog, but also the factors that pattern inter-individual aging variation among dogs. Specifically, we identified three key age-associated patterns in the methylome. First, we found that age exerts broad effects on DNAm across the genome, but that its impact is genome context dependent. Specifically, we found a broad age-associated hypermethylation of promoters that contrasted with age-associated loss of methylation in TEs. Second, X-linked LINE1 methylation was lower in females than males, illustrating potential differential susceptibility to age-associated decline. Third, dog size predicts the rate of age-associated decline in LINE1 DNAm, consistent with a potential role for LINE1 expression to explain a long-standing effect of dog size on aging rate and longevity. Collectively, these data deepen our understanding of the heterogeneous mammalian epigenome with relevance to health and age-associated phenotypes in dogs, and mammals more broadly.

Overall, age strongly patterned DNAm in companion dog immune cells, with the vast majority of age-associated loci exhibiting a loss of methylation with age–consistent with studies in other organisms [29,54]. However, the effect of age on DNAm varied across the genome and was highly context dependent. For instance, promoters were much more likely to exhibit age-associated gains in methylation and were enriched for numerous immune-response related and stress-responsive proteins and targets (**Supplemental Figure 3**). Age-related changes detected here may reflect age-associated dysregulation of key immune and regulatory genes (i.e. changes in the epigenetic accessibility of stress-responsive promoters and motifs likely would impact the ability for the organism to respond accordingly). In support of this, we found that promoters with age-associated hypermethylation were enriched for transcription factors involved in the stress responses. Methylation state of transcription factor binding sites has wide-ranging possible outcomes– but is generally associated with a decrease in transcription initiation [79–81]. One of these transcription factors, ATF4 (and its orthologs), is necessary for lifespan extension in important model organisms (e.g., *Saccharomyces cerevisiae*, *C. elegans*) and is also more abundant in tissues from long-lived mouse models [63–67]. ATF4 is also known to be involved in both viral and bacterial responses [62,82–84], so a change in methylation state of these sites could severely alter these immune cells’ ability to response to foreign nucleotides. Overall, these results support a loss of ability for older animals to respond to stressors–i.e., reduced resilience.

While promoters were largely hypermethylated with age, we found that most of the genome exhibited age-associated loss of DNAm, which was the strongest in TEs. Epigenetic dysregulation of TEs can contribute to aging, the development of age-associated diseases, and other disorders [36,46,85]. We also found that golden retrievers–a breed with a high incidence of hematopoietic cancer–show more rapid age-associated loss of methylation of LINE1s, which have been repeatedly linked to cancer [31,88] (**Figure 2F**). A genetic mechanism underlying this breed’s increased cancer risk has remained elusive, and those found only explain a fraction of the incidences [75,89]. While just one example, it raises the possibility of deploying reverse transcriptase inhibitors to delay or even prevent blood-based cancers in golden retrievers. The epigenetic accessibility and activity of TEs are broadly known to increase with age [32,46,86,87]. In mice, loss of *Sirt6*, a gene known to repress LINE1 transposition, can lead to progeria, which can be partially rescued by reverse-transcription inhibitor drug treatment [39]. Other recent work has uncovered that DNAm is lost at TEs, including LINE1s, with age in humans and non-human primates [32,54]. Our results significantly broaden our understanding of TEs and aging by capturing age-associated loss of LINE1 methylation in dogs. Thus we provide further support that loss of epigenetic repression of TE activity is a key feature of aging–though it is unclear whether increased TE activity is a cause or consequence of aging and age-related disease.

LINE1 elements are highly abundant on the X chromosome, which could be reflected in differences in LINE1 regulation between sexes. We found that, on average, X-linked LINE1s are more methylated in males compared to females, consistent with evidence in mammals that LINE1s aid in silencing in the context of X chromosome inactivation and are more epigenetically accessible on inactive X chromosomes [90–92]. It is unclear to what extent this difference in chromosome X’s methylation of LINE1s contributes to sex-specific aging, especially as non-DNAm modes of silencing may also control LINE1 expression. However, while not quantified here, it is possible that loss of epigenetic silencing of Y-linked LINE1s may be more damaging than X-linked LINE1s [93–95]. In support of this, studies of mice have found that genetically modified XY females lived shorter than normal XX females, and genetically modified XX males lived longer than XY males [96]. Human aneuploidy studies have also found XYY humans have life expectancy that is 10 years shorter than XY humans, while this reduction in life expectancy relative to XY is only 2 years in XXY humans [97,98].

Our results suggest that: (i) dog size recapitulates age-associated DNAm patterns such that larger dogs have “older” epigenomes than their similarly-aged counterparts, (ii) age-associated X-linked LINE1’s are more highly methylated in males than females, and (iii) large dogs show the fastest age-associated hypomethylation in LINE1 elements (**Figure 4**). Researchers commonly propose that differences in exposure to growth hormone and differential insulin-like growth factor signaling explain size-related lifespan variations and the onset of age-related diseases within a species, including studies in humans [99–102]. This is in light of extensive and foundational work done in model organisms, including *S. cerevisiae, Drosophila,* and *C. elegans,* tying insulin-like growth-factor signaling to the progression of aging [1,101,103–108]. Compelling studies suggest that exposure to insulin-like growth factor signaling, especially during development, contributes to differences in dog life expectancy [109,110]. Our data presented here offers an additional perspective that TEs may also be impacting life expectancy and disease states across sizes in addition as well.

Although these data do not quantify the degree to which LINE1s are transposing with age and are cross-sectional in nature, these findings broadly impact aging studies, especially given recent findings that reverse-transcriptase inhibitors can delay aging and diseases of aging by suppressing transposable elements [40,41,111].

Could the differences in LINE1 epigenetic regulation (e.g. DNAm) found here be reflected in other contexts important to lifespan? Other differences in demographics within dogs are important in the progression of lifespan (e.g. sterilization status [112], social [5,113], environmental [114]). Understanding the epigenetic differences under circumstances that influence lifespan may lead us to a common mechanism responsible for the differences in aging and disease onset within a species. The DAP has already enlisted more than 40,000 dogs that are intriguingly from various genetic and environmental backgrounds. The 864 dogs studied here belong to a cohort of ∼1,000 dogs, the “Precision Cohort,” that are to be studied longitudinally in the Dog Aging Project, collecting blood samples and sequencing the epigenome yearly in the same manner as described here. This project will continue to uncover the genetic, metabolomic, and epigenetic prospects contributing to within-species differences in aging. It will also uncover to what extent LINE1s are differentially regulated across different demographics and lead to a general understanding of how they may more broadly influence dog disease states and lifespan itself.

## Methods

### Blood sample collection

This study examined changes in DNA methylation (DNAm) in peripheral blood mononuclear cells (PBMCs) from whole blood samples collected annually from the Precision cohort, starting in 2021 up until February 2024. Dogs were enrolled in the Precision cohort on a rolling basis, with veterinarians collecting samples annually based on each dog’s enrollment date (**Figure 1A**). These blood samples were refrigerated and sent to the DAP team members at Texas A&M University, which are then aliquotted and sent on ice to the Arizona State University (ASU) DAP team. The ASU DAP team isolated peripheral blood mononuclear cells (PBMCs) from the whole blood samples [115]. PBMCs offer significant advantages for analysis as (1) they are easily attainable from a minimally invasive blood draw, (2) they consist of a diverse set of cells (lymphocytes, monocytes, and macrophages), and (3) their methylation patterns have been found representative to human health and disease [27,116,117].

### Sample annotation

The paired metadata for this study were drawn from the Dog Aging Project’s Health and Life Experience Survey (HLES), which collects extensive information about dogs’ activity, environment, behavior, and general characteristics. For this study, breed background data from HLES were particularly relevant, as they enabled the assignment of dogs to one of five size categories: Toy and Small, Medium, Standard, Large, and Giant [44].

To further refine the analysis, predicted adult heights for the dogs were calculated using a machine learning model based on DAP height survey data, with height values ranging from 0 to 4 (0: ankle high; 1: calf high; 2: knee high; 3: thigh high; 4: waist high). This model was trained using genetic information from thousands of dogs, as described previously [78]. The predicted heights were strongly correlated with the traditional breed size categories (**Supplemental Figure 1B**).

Using predicted heights as a proxy for breed size offers several advantages: (1) it reduces subjectivity in size classification by positioning dog breed size on a continuous scale, (2) it accounts for pre-adult dogs by consistently reflecting predicted adult size throughout the dog’s lifespan, allowing for the inclusion of dogs that are not fully grown, and (3) it captures the inherent non-linearity of breed size categories, offering a more nuanced representation of size variation among dogs.

The University of Washington IRB deemed that recruitment of dog owners for the Dog Aging Project, and the administration and content of the DAP Health and Life Experience Survey (HLES), are human subjects research that qualifies for Category 2 exempt status (IRB ID no. 5988, effective 10/30/2018). All study-related procedures involving privately owned dogs were approved by the Texas A&M University IACUC, under AUP 2021-0316 CAM (effective 12/14/2021).

### RRBS preparation, sequencing, and data preprocessing

DNA was then extracted for reduced-representation bisulfite sequencing (RRBS) using the Quick-DNA/RNA Magbead kit (Zymo Research, Irvine, California, USA) [115]. The Translational Genomics Research Institute (Phoenix, Arizona) sequenced the samples on the Illumina NovaSeq S4 platform in collaboration with the ASU DAP team members.

We trimmed reads with Trim Galore! and aligned the reads to the NCBI UU_Cfam_GSD_1.0 (CanFam4) reference genome [118]. To extract methylation and read count data, we used Bismark [119]. We removed samples with less than one million mapped reads and re-sequenced these samples if the sample amounts allowed.

We kept CpG sites with at least 5x coverage in at least 33% of the samples for downstream analysis [50]. CpG sites that were consistently methylated (>90% mean percent methylation) and consistently unmethylated (<10% mean percent methylation) in samples were also removed. We expect neighboring CpG sites to have similar DNA methylation patterns, given their relative proximity in the genome (as shown by the literature), so we grouped these CpG sites into regions using BSseq [51,120]. In our framework, we required CpG sites to be closer than 250 base pairs to be grouped into the same region and the region to have at least 5x coverage in 33% of the samples (**Supplemental Figure 2A**). In total, we generated 47,393 regions for downstream analyses. These regions vary in length, but most have, on average, ∼25 nucleotides between individual CpG sites (**Supplemental Figure 2B**). Regions that were almost always methylated (>90% percent methylation) or unmethylated (<10% mean percent methylation) across the samples were removed so that we could test the variably methylated regions (**Supplemental Figure 2C**). ∼80% of all regions had 5x or more region coverage across 95% of all 864 samples (**Supplemental Figure 2D**). Moreover, ∼80% of all samples had 5x or more coverage across 95% of all 47,393 regions (**Supplemental Figure 2E**).

### Region annotation

We obtained gene information from the NIH CanFam4 gene annotation file. We defined promoters as regions 2kb upstream of transcriptional start sites, and when annotation was available, we identified introns. We downloaded the CpG islands and TEs locations from the CanFam4 UCSC genome browser [121]. We generated annotations for CpG shores and CpG shelves in R, where CpG shores are the 2kb that flank CpG Islands and CpG shelves flank CpG shores by 2kb [122]. We annotated regions overlapping chromatin state annotations (e.g. heterochromatin, quiescent) using Son et al.’s mapping of the dog epigenome [123].

### Region-based differential methylation analysis

PQLSeq, a binomial mixed modeling framework, was used to model differential methylation as a function of age, while controlling for sex, predicted height, and genetic relatedness across 864 samples [51,124]. Functional class scoring used the R package fgsea with the provided curated gene sets [125,126].

### Instruments and statistical methods

All bioinformatics analyses were conducted on the Sol Supercomputer at ASU [127] and with R (v4.2.2). We corrected for multiple hypothesis tests using Benjamini-Hochberg’s False Discovery Rate (FDR). Estimates used linear models accounting for age, with code provided upon peer-reviewed publication.

### Data sharing and availability

The Dog Aging Project is an open data project. These data are housed on the Terra platform at the Broad Institute of MIT and Harvard. Interested researchers and the general public can apply for data access at dogagingproject.org/open_data_access.

## Acknowledgments

The authors thank Dog Aging Project participants, their dogs, and community veterinarians for their essential contributions. This research is based on publicly available data collected by the Dog Aging Project, under U19 grant AG057377 (PI: Daniel Promislow) from the National Institute on Aging, a part of the National Institutes of Health, and by additional grants and private donations, including generous support from the Glenn Foundation for Medical Research, the Tiny Foundation Fund at Myriad Canada, the WoodNext Foundation, and the Dog Aging Institute. These data are housed on the Terra platform at the Broad Institute of MIT and Harvard. BMM is supported by an NIH F99/K00 and a Graduate Research Fellowship from the National Science Foundation. DP received support from USDA cooperative agreement USDA/ARS 58-8050-9-004. The authors acknowledge Research Computing at Arizona State University for providing High Performance Computing and Storage resources that have contributed to the research results reported within this paper. The content is solely the authors’ responsibility and does not necessarily represent the official views of the National Institutes of Health.

## Dog Aging Project Consortium Authors (as of March 2022)

Joshua M. Akey^1^, Brooke Benton^2^, Elhanan Borenstein^3^, Marta G. Castelhano^4^, Amanda E. Coleman^5^, Kate E. Creevy^6^, Kyle Crowder^7^, Matthew D. Dunbar^8^, Virginia R. Fajt^9^, Annette L. Fitzpatrick^10^, Unity Jeffery^11^, Erica C Jonlin^12^, Matt Kaeberlein^13^, Elinor K. Karlsson^14^, Kathleen F. Kerr^15^, Jonathan M. Levine^16^, Jing Ma^17^, Robyn L McClelland^18^, Daniel E.L. Promislow^19^, Audrey Ruple^20^, Stephen M. Schwartz^21^, Sandi Shrager^22^, Noah Snyder-Mackler^23^, Katherine Tolbert^24^, Silvan R. Urfer^25^, Benjamin S. Wilfond^26^

^1^ Lewis-Sigler Institute for Integrative Genomics, Princeton University, Princeton, NJ, USA

^2^ Department of Laboratory Medicine and Pathology, University of Washington School of Medicine, Seattle, WA, USA

^3^ Department of Clinical Microbiology and Immunology, Sackler Faculty of Medicine, Tel Aviv University, Tel Aviv, Israel

^4^ Cornell Veterinary Biobank, College of Veterinary Medicine, Cornell University, Ithaca, NY, USA

^5^ Department of Small Animal Medicine and Surgery, College of Veterinary Medicine, University of Georgia, Athens, GA, USA

^6^ Department of Small Animal Clinical Sciences, Texas A&M University College of Veterinary Medicine & Biomedical Sciences, College Station, TX, USA

^7^ Department of Sociology, University of Washington, Seattle, WA, USA

^8^ Center for Studies in Demography and Ecology, University of Washington, Seattle, WA, USA

^9^ Department of Veterinary Physiology and Pharmacology, Texas A&M University College of Veterinary Medicine & Biomedical Sciences, College Station, TX, USA

^10^ Department of Family Medicine, University of Washington, Seattle, WA, USA

^11^ Department of Veterinary Pathobiology, Texas A&M University College of Veterinary Medicine & Biomedical Sciences, College Station, TX, USA

^12^ Department of Laboratory Medicine and Pathology, University of Washington School of Medicine, Seattle, WA, USA

^13^ Department of Laboratory Medicine and Pathology, University of Washington School of Medicine, Seattle, WA, USA

^14^ Bioinformatics and Integrative Biology, University of Massachusetts Chan Medical School, Worcester, MA, USA

^15^ Department of Biostatistics, University of Washington, Seattle, WA, USA

^16^ Department of Small Animal Clinical Sciences, Texas A&M University College of Veterinary Medicine & Biomedical Sciences, College Station, TX, USA

^17^ Division of Public Health Sciences, Fred Hutchinson Cancer Research Center, Seattle, WA, USA

^18^ Department of Biostatistics, University of Washington, Seattle, WA, USA

^19^ Department of Laboratory Medicine and Pathology, University of Washington School of Medicine, Seattle, WA, USA

^20^ Department of Population Health Sciences, Virginia-Maryland College of Veterinary Medicine, Virginia Tech, Blacksburg, VA, USA

^21^ Epidemiology Program, Fred Hutchinson Cancer Research Center, Seattle, WA, USA

^22^ Collaborative Health Studies Coordinating Center, Department of Biostatistics, University of Washington, Seattle, WA, USA

^23^ School of Life Sciences, Arizona State University, Tempe, AZ, USA

^24^ Department of Small Animal Clinical Sciences, Texas A&M University College of Veterinary Medicine & Biomedical Sciences, College Station, TX, USA

^25^ Department of Laboratory Medicine and Pathology, University of Washington School of Medicine, Seattle, WA, USA

^26^ Treuman Katz Center for Pediatric Bioethics, Seattle Children’s Research Institute, Seattle, WA, USA

## Supplementary figures

**Supplemental Figure 1:**
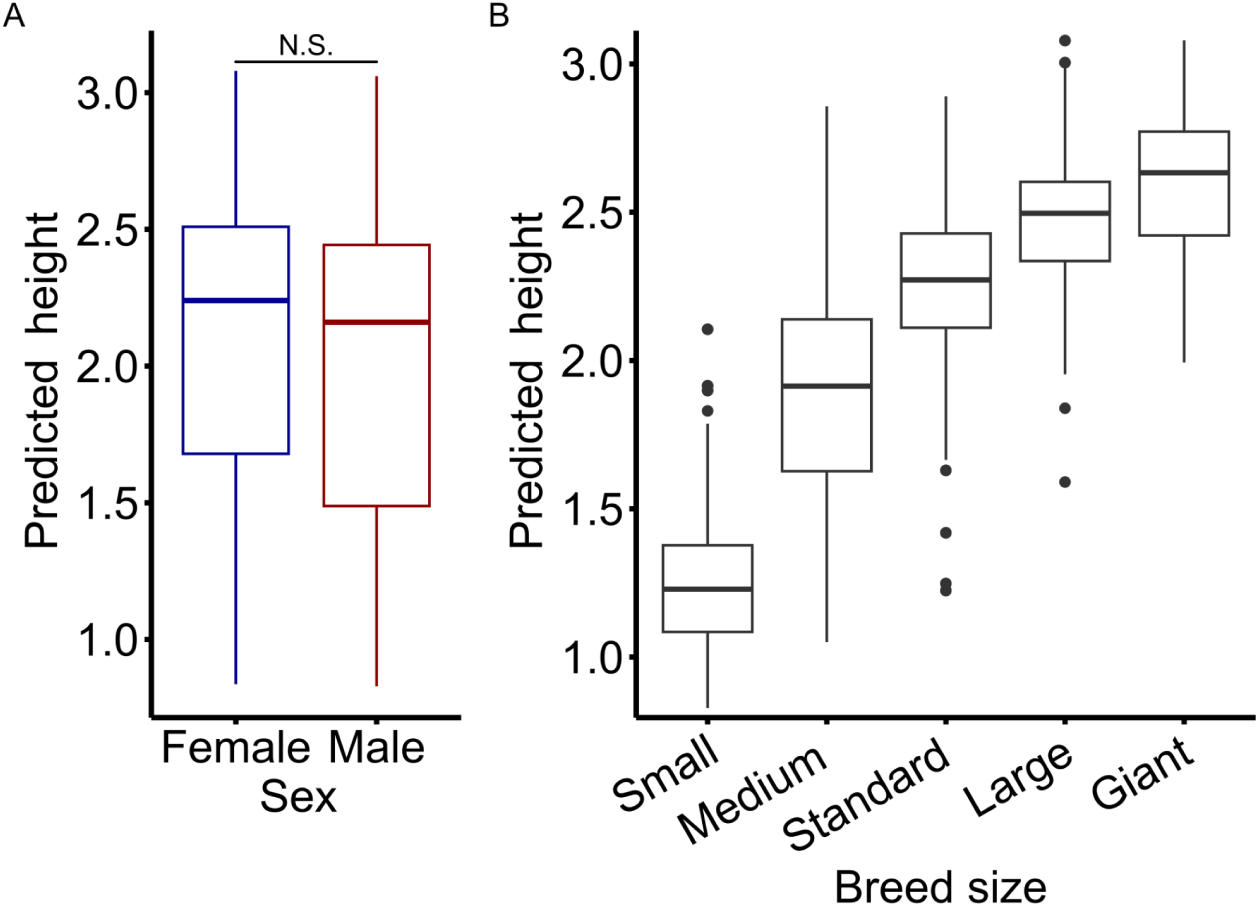
Breed size and predicted height overview. A) In this study, the dog size (predicted height) of female dogs and the dog size of the male dogs (Student’s t-test, p = 0.12). B) Dog size (predicted adult height) versus breed size class of dogs in the study.

**Supplemental Figure 2:**
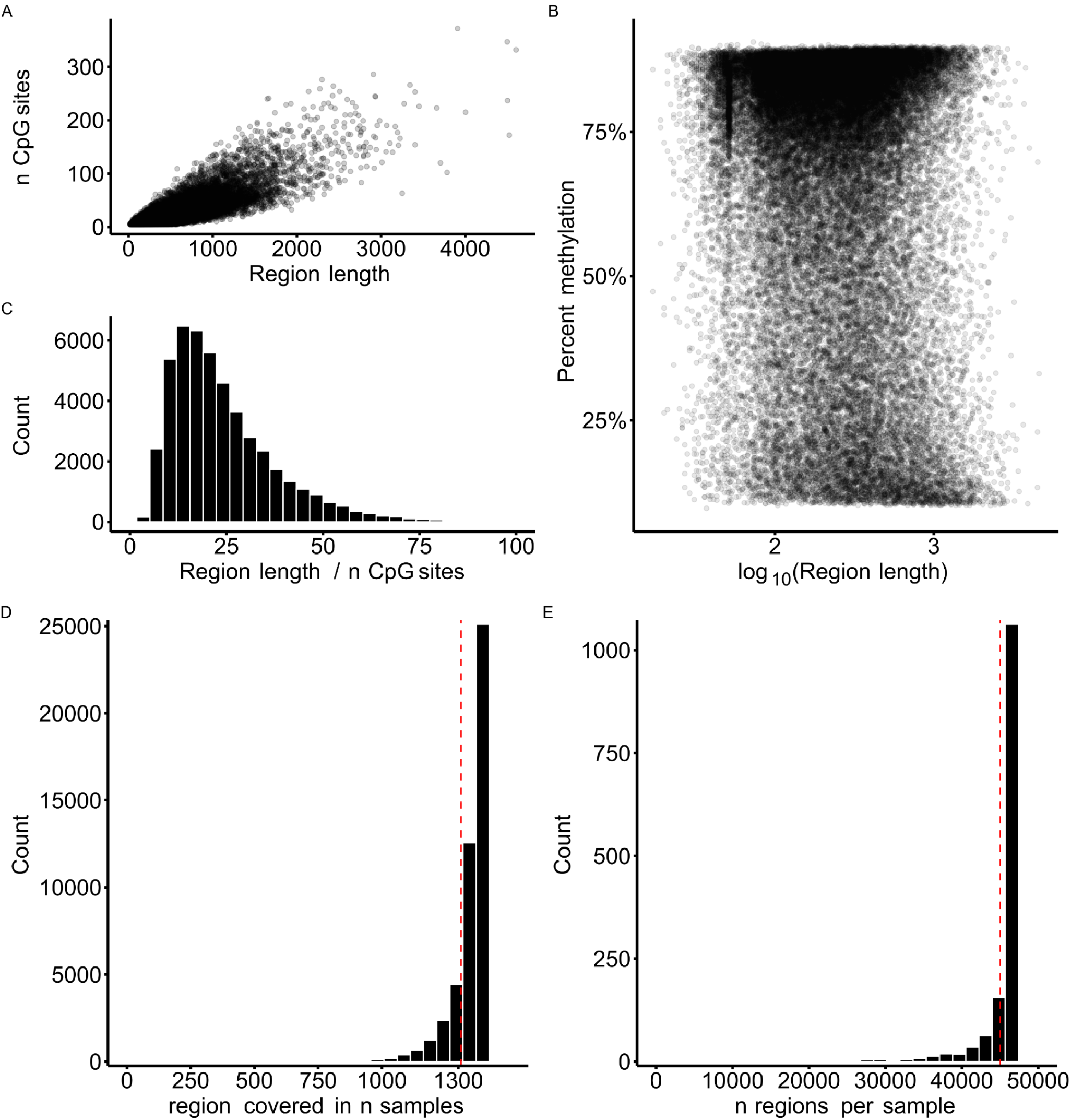
Sample and region overview. A) Number of CpGs over the length of the 47,393 regions used in this study. B) Region length / CpG sites for all regions used in this study. C) Distribution of mean percent methylation and region length of the regions across the 864 samples. D) Most regions had 5x or more coverage across 95% of samples (95% threshold shown in red). E) Most samples had 5x coverage of 95% of regions (95% threshold shown in red).

**Supplemental Figure 3:**
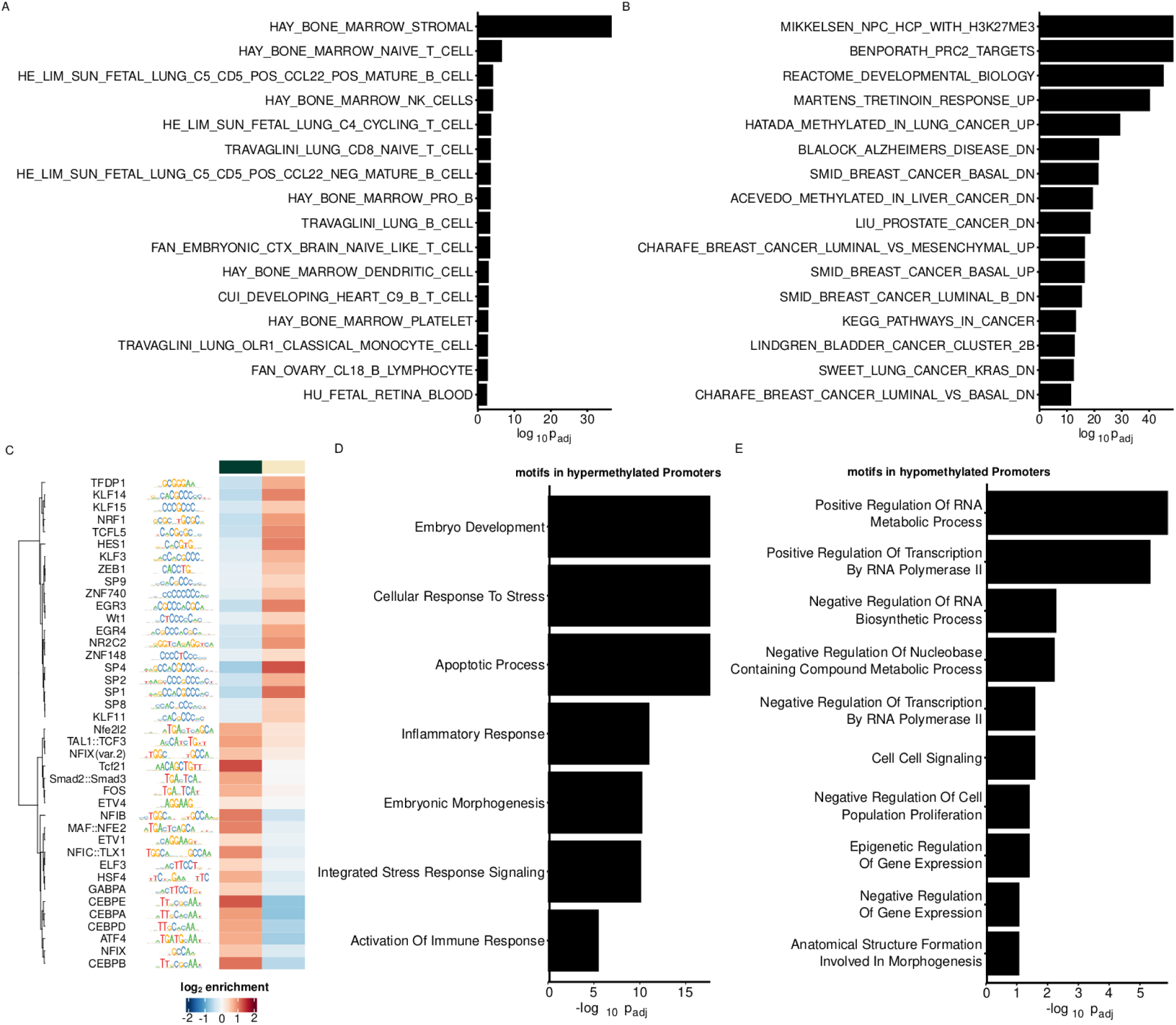
Promoter set and transcription factor binding site enrichment. Functional class enrichment using fgsea functional class scoring (methods) of fgsea provided (A) cell type signature gene set and (B) curated gene set ranked by standardized effect of age [125]. C) Motif enrichment results among top 20 promoter regions hypomethylated and hypermethylated with age, all with FDR < 0.01 [128]. At the top of the heatmap, green indicates the regions hypermethylated with age, while tan indicates the regions hypomethylated with age. D, E) Ontology enrichment of proteins whose promoter targets are (D) hypermethylated and (E) hypomethylated with age.

